# Effects of high fat and sugar diet on motivation for food and resistance to punishment in rats: role of sex and age of exposure

**DOI:** 10.1101/2023.11.13.566808

**Authors:** Stevenson Desmercieres, Virginie Lardeux, Jean-Emmanuel Longueville, Emilie Dugast, Nathalie Thiriet, Marcello Solinas

## Abstract

Exposure to food rich in fat and sugar (High Fat and Sugar Diet, HFSD) is believed to induce behavioral and neurobiological changes that would produce addiction-like behavior and increase the risks of obesity and overweight. Studies in rodents have led to conflicting results suggesting that several factors such as sex and age of exposure contribute to the development of maladaptive behavior towards food. In addition, it is not clear whether the effects of exposure to HFSD persist after its discontinuation which would indicate long-term risk to develop addiction-like behavior. In this study, we investigated the persistent effects of an intermittent 8-week exposure to HFSD in male and female rats as a function of age of exposure (adult and adolescent). We found that intermittent exposure to HFSD did not alter body weight, but it affected consumption of standard food during the time of exposure in all groups. In addition, in adults, HFSD produced a decrease in the initial baseline responding in FR1 schedules that persisted for 4 weeks in males but not in female rats. However, we found that exposure to HFSD did not affect resistance to punishment measured by progressive shock strength (PSS) break points or motivation for food measured by progressive-ratio break points regardless of sex or age of exposure. Altogether, these results do not provide support to the hypothesis that intermittent exposure to HFSD produce persistent increases in the vulnerability to develop addiction-like behaviors towards palatable food.

## INTRODUCTION

The prevalence of obesity and overweight has dramatically increased in the world in the last decades (World Health Organization, www.who.int). Although obesity has increased in the entire population, the increases appear more pronounced among children, adolescents and women (WHO, www.who.int) suggesting that development and hormonal factors can have an impact on this condition. Importantly, obesity is associated with increased mortality, morbidity and impaired quality of life (Abdelaal et al., 2017) and, an analysis of societal costs has estimated that its economic impact is on average 1.8% of gross domestic product and may increase up to 3.6% in 2060 (Okunogbe et al., 2021). Therefore, understanding the factors that influence obesity and the underlying biological mechanisms is critical to design strategies to limit its spreading.

In humans, not only continuous consumption, but also intermittent access to food rich in fat and sugar, separated by self-imposed periods of dieting, contributes to the development of obesity and forms of eating disorders (Dulloo et al., 2015). These experiences of repeated cycles of highly palatable food consumption and diet with ‘safe food’ could alter the control of feeding (Dulloo et al., 2015). In rats, several studies have shown that an intermittent access to highly palatable food, typically composed by a high concentration of fat and sugar (High Fat and Sugar Diet, HFSD), with standard chow diet promote behaviors similar to those found with abuse drugs. For example, intermittent access to HFSD produces increases of compulsive behavior in rats (Rossetti et al., 2014), withdrawal-like states characterized by the emergence negative emotional state and anxiety (Cottone et al., 2009; Iemolo et al., 2012), decreases in reward system functioning (Moore et al., 2020) and impairments of memory function and hippocampal neurogenesis (Ferragud et al., 2020). Nevertheless, the long-lasting consequences of exposure to HFSD on behavior remains unclear and preclinical studies are essential to determine the behavioral, physiological and neurobiological mechanisms underlying the development of maladaptive behavior as food disorders, that lead to obesity (O’Connor and Kenny, 2022).

Two main measures are often used to characterize the addiction-like behavior induced by HFSD: excessive motivation measured by progressive ratio (PR) schedules (Hodos, 1961) and compulsivity measured by resistance to punishment (George et al., 2022; Vanderschuren and Ahmed, 2013). Several studies have investigated the effects of exposure to HFSD on motivation and compulsivity and have obtained conflicting results. For example, some studies found increases in motivation (Wojnicki et al. 2006; la Fleur et al. 2007; Figlewicz et al. 2013; Garman et al. 2021), some found decreases (Davis et al. 2008; Vendruscolo et al. 2010; Blaisdell et al. 2014; Tracy et al. 2015; Wong et al. 2017) and others found no effect (Jong et al. 2013; Rossetti et al. 2014; Tracy et al. 2015). Similarly, for resistance to punishment, some studies found increases (Johnson and Kenny 2010; Oswald et al. 2011; Rossetti et al. 2014) and one found no effect (Jong et al. 2013). These discrepancies may be due to differences in the sex and the age of the animals, duration and type of exposure, eventual discontinuation of treatment and the PR or punishment procedures used.

We have recently developed a self-adjusting procedure to investigate punishment in rats (Desmercieres et al., 2022). The progressive shock strength (PSS), similar to procedures commonly used in humans (Apergis-Schoute et al., 2017; Kanen et al., 2021; Kim and Anderson, 2020), allows individuals to titrate the level of punishment they are willing to receive in order to obtain a reward and provides PSS breakpoints that are sensitive to the motivation state and the value of the reward (Desmercieres et al., 2022). This procedure allows a continuous rather than dichotomic evaluation of resistance to punishment and may allow to more precisely measure the consequence of exposure to HFSD on resistance to punishment.

In this study, we used the PR and the PSS procedures to assess the effects of HFSD on motivation and compulsivity for sweet rewards. Because sex is an important risk factor for severe obesity (Hales, 2020) and eating disorders, we investigated the effects of HFSD in both male and female rats. In addition, because adolescence is a critical period for brain development (Spear, 2000) and a window of vulnerability for the development of maladaptive behaviors including addiction (Chambers et al., 2003) and eating disorders (Klump, 2013), we investigated the effects of exposure to HFSD in both adolescence and adult rats.

## METHODS

### Subjects

Sprague-Dawley (Janviers Labs) rats of both sexes were used: 47 adolescents (4 weeks of age at the beginning of the HFSD exposure; 155 – 160 g for females and 130 – 135 g for males) and 48 adults (10 weeks of age at the beginning of the HFSD exposure; 230 – 250 g for females and 320 – 350 g for males). All rats, experimentally naïve at the start of the study, were housed in pairs with two wooden sticks for enrichment, in individually ventilated cages (Techniplast, Sealsafe Plus GR900), in a temperature- and humidity-controlled room kept under reversed light-dark cycle conditions (12 hours light-dark cycle, lights off at 7 AM). All experiments were conducted during the dark phase and in accordance with European Union directives (2010/63/EU) for the care of laboratory animals and approved by the local ethics committees (COMETHEA).

### General design of the experiment

General design of the experiment is illustrated in fig. 1. First, rats were exposed to chronic HFSD or control diet for 8 weeks. At the end of this period, HFSD was discontinued and standard diet was used for all groups until the end of the experiment. Food self-administration started one week after the last exposure to HFSD and Mild food restriction (∼15 g/day for females and ∼20 g/day for males) was implemented to motivate food responding and limit weight gain and maintain stable behavior until the end of the experiment. Food was given 1 hour after the end of the experimental sessions. Rats had unlimited access to water. Rats were allowed to self-administer food pellets for 9 training sessions and then resistance to punishment and food motivation were measured alternatively once per week for 6 weeks. A typical week included 4 training and 1 test session. At the end of the experiment, anxiety levels and sensitivity to pain were measured on the same day.

**Fig 1:**
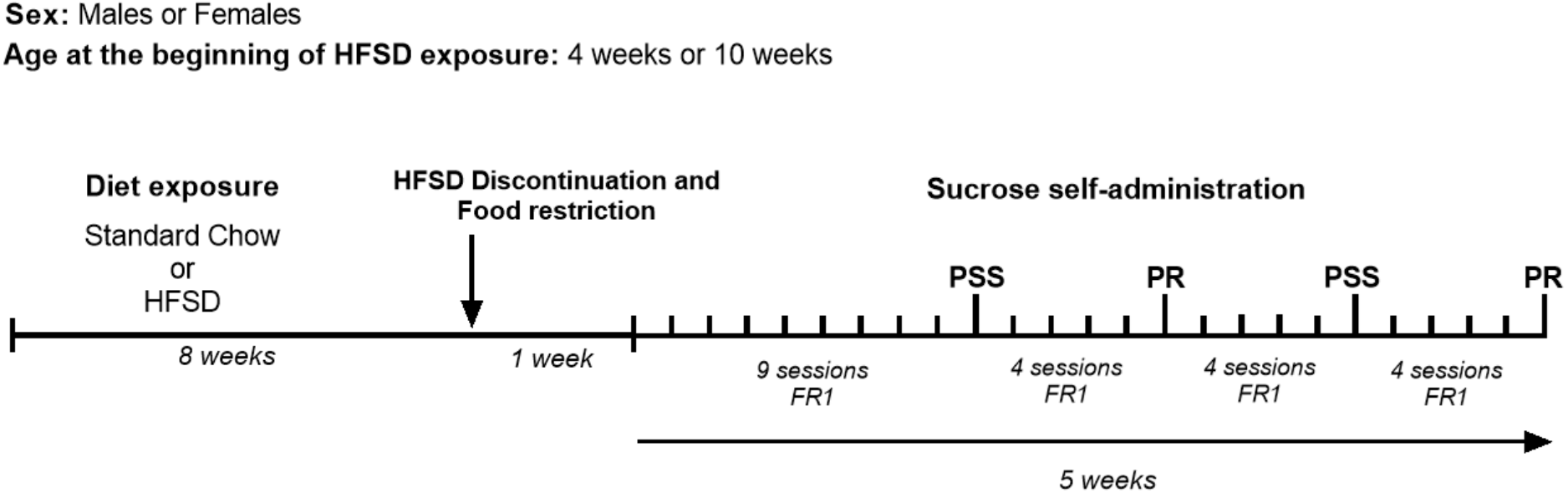
General Experimental Design. Male or female rats were exposed to HFSD or control diet starting at adolescence (4 weeks of age) or early adulthood (10 weeks of age) for 8 weeks according to an intermittent schedule in which HFSD was only available on weekends whereas during the other weekdays all rats had only access to standard chow. After 1 week of discontinuation of HFSD and mild food restriction, sucrose self-administration started with a FR1 schedule for 9 training sessions and then PSS and PR sessions were conducted every fifth day (on Friday) alternatively with interspersed additional FR1 sessions. Note that for adolescent groups, exposure to HFSD started during adolescence and was maintained until young adulthood when sucrose self-administration was performed.

We performed 4 separate experiments: 1) effects of HFSD in adolescent males; 2) effects of HFD in adolescent females; 3) effects of HFSD in adult males; and 4) effects of HFSD in adult females.

#### Exposure to HFSD

We used the HFSD exposure procedure described by Rossetti et al., which was shown to induce compulsivity for food in female rats (Rossetti et al., 2014). Briefly, half of the rats were assigned to HFSD condition and the control condition. HFSD rats had access to 5TCY Tablets (Test diets, LabTreat^TM^ OmniTreat^TM^ Enrichment Tablet, 5g/Tablet) for the weekends (Friday 6:00 p.m. to Monday 9:00 a.m.) and the rest of the week, they had access to standard food (4RF21, Mucedola, Italy) in their homecage for eight weeks. Control rats had access only to standard food (4RF21, Mucedola, Italy) for the same period of time.

HFSD and standard diets had a similar number of calories (see Table. 1) and were given *ad libitum* during the exposure.

**Table 1:** Diet composition.

| Food | Energy density (kcal/g) | Macronutrient composition (kcal%) |  |  |
| --- | --- | --- | --- | --- |
|  |  | Carbohydrate | Protein | Fat |
| 4RF21 Mucedola (standard food) | 3.150 | 46.31 | 18.5 | 2.6 |
| 5TCY Test diets (High Fat Sugar Diet) | 3.290 | 63.6 | 22.5 | 13.9 |
| Precision pellet 5TUT 45 mg (sweet pellets) | 3.404 | 100 | 0 | 0 |

### Food Reinforcement Apparatus and training procedure

Experimental chambers (MedAssociates, www.medassociates.com) were enclosed individually in sound-attenuation chests. Each experimental chamber had a recessed food tray, and two levers in the right wall. The floor was constituted by bars that were connected to shockers that could deliver foot shock, with electric current set to 0.45 mA. Each chamber was equipped with a food-pellet dispenser, which could deliver 45 mg pellets to the food tray. Experimental events were controlled by computers using MedAssociates interface and Med-PC IV software; Med-PC code used to conduct the procedures is available upon request. A diode light was present on each lever. One lever was assigned to be the active lever and the corresponding light was used as a conditioned stimulus for food reinforcement. A third diode light was installed on the opposite wall, and its flashing was used as a discriminative stimulus to indicate that food reinforcement would be associated with a foot shock.

The general training consisted of 45-min sessions of a fixed-ratio 1 (FR1) schedule of food reinforcement in which one lever press was required to obtain a 45 mg sucrose pellet. During these sessions, food availability was signaled by turning off the house-light, indicating that a single response on the active lever would immediately result in the delivery of one food pellet, accompanied by flashing of the diode light above the lever for 2 secs. Subsequently, the house light was turned on for an additional 18-sec time-out period, during which responding had no programmed consequences. Following the time-out, a new trial started and the next response on the right lever was again reinforced. Responses on the inactive lever were recorded but never reinforced.

### Self-adjusting progressive shock strength (PSS) procedure

Resistance to punishment was measured in the 10^th^ and 20^th^ session as previously described (Desmercieres et al., 2022). In the PSS procedure, active lever presses resulted in the delivery of food rewards and foot-shocks of different strengths. The PSS consisted of steps in which the shock duration was increased each time the animal obtained 2 separate rewards at a given duration. The duration of the first step was 0 sec (no shock), the second step was a low duration of 0.05 sec and subsequent shocks increased of 15% at each step for 20 steps. Thus, the step sequence of durations was: 0, 0.05, 0.06, 0.07, 0.08, 0.09, 0.10, 0.12, 0.13, 0.15, 0.18, 0.20, 0.23, 0.27, 0.31, 0.35, 0.41, 0.47, 0.54, 0.62, and 0.71 sec. At the beginning of each shock trial, the light on the opposite side of the levers and food tray, was switched on and off intermittently for the entire trial for periods of time proportional to the duration of the shock to signal the presence of shock contingencies according to the formulas: Duration On = 0.1 sec times step number and Duration Off = 2 sec – Duration On. If animals reached the final step, the duration of the shock was not further increased, and all subsequent shocks were set at 0.71 sec. If rats did not emit any response for 5 min, shock duration was reset to 0 sec and the shock progression was reinitialized. The 5 min threshold for reset was chosen based on pilot experiments showing that this level allowed most animals to reinitiate their operant behavior within a single session. Thus, animals could at any moment avoid higher strength of shock by limiting the frequency of food reinforcement until shock duration returned to 0 sec, which was signaled by the absence of the blinking light. The strength of the shock was measured by the electrical charge in millicoulombs (mC) that an animal would tolerate to obtain food pellets and was calculated by multiplying the fixed current of the shock (0.45 mA) by the duration in sec. PSS sessions lasted 45 minutes and the break point was calculated by the cumulated electrical charge sustained during the session.

### Progressive-ratio (PR) schedule

Animal motivation was measured in the 15^th^ and 25^th^ session. Under the PR schedule of food reinforcement, the number of responses required to obtain a food pellet increased with each successive food pellet. The steps of the exponential progression were the same as those previously developed by Roberts and colleagues (Richardson and Roberts, 1996) adapted for food reinforcement (Solinas et al., 2003; Solinas and Goldberg, 2005), based on the equation: response ratio = (5e^(0.2^ ^×^ ^reinforcer^ ^number)^) − 5, rounded to the nearest integer. Thus, the values of the steps were 1, 2, 4, 6, 9, 12, 15, 20, 25, 32, 40, 50, 62, 77, 95, 118, 145, 178, 219, 268, 328, 402, 492, 603, and 737. Sessions under the progressive-ratio schedule lasted until 10 min passed without completing a step, which, under basal conditions, occurred within 1 hour.

### Anxiety-related behaviors: Elevated plus maze procedure

The elevated plus maze (Viewpoint, Lyon, France) was composed of 4 arms (50 cm long * 10 cm wide at 42,5cm from the floor). Two arms had no walls (considered as open arms) and two had black walls on its edges (considered as closed arms). Each arm was placed on the opposite side of its copy separated by a central square (10*10 cm) in the middle giving access to all the arms. Rats position was recorded automatically and in real time by a camera and video tracking software (Viewpoint, Lyon, France). The software defined 4 virtual areas delimiting with a center zone. Anxiety was measured as the time spent in open arms so that more time spent in open arms indicated a lower level of anxiety.

### Pain sensitivity: Hot-plate test

The hot-plate (Ugo Basile, model-DS 37) was maintained at 48 °C (Deuis et al. 2017). After a habituation of 10 min, animals were placed into a glass cylinder of 25 cm diameter on the heated surface and 47 centimeters walls. The latency before escape or jumping, was recorded. Experiments were stopped after a cut-off of 120 sec to prevent unnecessary pain or tissue damage.

### Statistical Analysis

Data were analyzed in GraphPad Prism. Data were checked for normality distribution using the Shapiro–Wilk test. When the assumption of sphericity had been violated, data were analyzed using analysis of variance (ANOVA) corrected by the Greenhouse–Geisser method. Data Weight gain and food consumption were assessed by two-way ANOVA for repeated measures followed by Sidak’s multiple comparisons post hoc test. PSS data were normalized using natural logarithm transformation as previously done (Desmercieres et al., 2022). For baseline measures of PSS and PR break points, we used two-way ANOVA for repeated measures with time and diet as factors followed by Sidak’s multiple comparisons post hoc test. Anxiety level and pain sensibility were assessed with a Mann-Whitney test because Shapiro-Wilk test revealed that distribution of data was not normal. Differences were considered significant when P < 0.05.

## RESULTS

### HFSD does not alter body weight but alters food consumption during the period of exposure

Each HFSD group begins a similar weight and increase similarly during the diet exposure compared to their control group. In adult male rats, before exposure to HFSD, the control group weight 359 ± 4 g and 355 ± 3 g for the HFSD group (Fig. 2A). 8 weeks later, at the end of the experiment, both groups showed similar weights (610 ± 8 g for the control group and 618 ± 9 g for the HFSD group) (Fig. 2A). Statistical analysis on body weight revealed a significant effect of time (F (1.584, 34.85) = 1966, P<0.0001; Greenhouse–Geisser ε = 0.20), no effect of diet (F (1, 22) = 0.03016, P=0.8637) and no significant time X diet interaction (F (8, 176) = 0.9746, P=0.4574). In adult female rats, the control group weighted 243 ± 3 g and the HFSD group weighted 244 ± 3 g before diet exposure (Fig. 2B). After 8 weeks, both groups showed similar weights (309 ± 5 g for the control group and 311 ± 4 g the HFSD group) (Fig. 2B). Statistical analysis on body weight revealed a significant effect of time (F (3.448, 75.86) = 433.6, P<0.0001; Greenhouse–Geisser ε = 0.43), no effect of diet (F (1, 22) = 0.4023, P=0.5325) and no significant time X diet interaction (F (8, 176) = 0.4958, P=0.8581). In adolescent male rats, initially HFSD and controls weighted 136 ± 2 g and 130 ± 2 g respectively (Fig. 2C). After 8 weeks, both groups showed similar weights (547 ± 6 g for the control group and 544 ± 7 g for the HFSD group) (Fig. 2C). Statistical analysis on revealed a significant effect of time (F (2.295, 50.50) = 7326, P<0.0001; Greenhouse–Geisser ε = 0.29), no effect of diet (F (1, 22) = 0.4127, P=0.5272) but a significant time X diet interaction (F (8, 176) = 1.993, P=0.0498). In adolescent female rats initially HFSD and controls weighted 156 ± 3 g and 159 ± 2 g respectively (Fig. 2D). After 8 weeks, both groups showed similar weights (304 ± 7 g for the control group and 307 ± 4 g for the HFSD group) (Fig. 2D). Statistical analysis on body weight revealed a significant effect of time (F (2.887, 60.64) = 499.4, P<0.0001; Greenhouse–Geisser ε = 0.36), no effect of diet (F (1, 21) = 0.1359, P=0.7161) and no significant time X diet interaction (F (8, 168) = 0.8958, P=0.5214).

**Fig 2:**
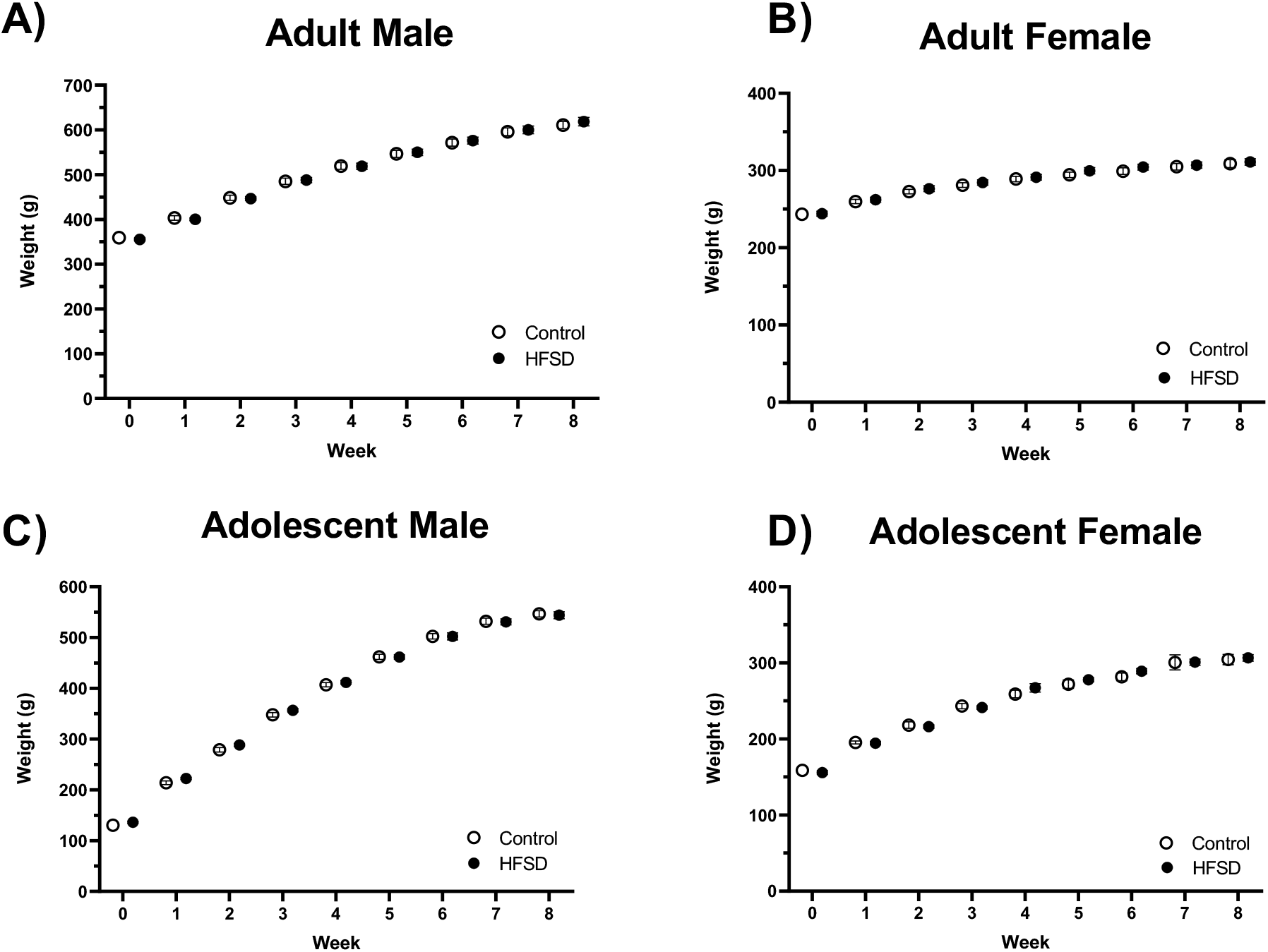
Effects of HFSD on body weight. Time course of body weight during the 8 weeks of exposure to HFSD in rats exposed to standard food (Control) and rats exposed to HFSD (HFSD). A) Adult males (HFSD n=12 ; Control n=12); B) Adult females (HFSD n=12 ; Control n=12); C) Adolescent males (HFSD n=12 ; Control n=12) and D) Adolescent females (HFSD n=12 ; Control n=11). Data are expressed as Mean ± SEM.

During the weekends, when HFSD group had access to fat and sugar food and control group to standard food, some differences on the food consumption can be relieved. In adult male rats, food consumption was similar in the two groups except on the first (30.5 ± 0.2 g/day for the control group compared to 29.1 ± 0.2 g/day for the HFSD group) and third weekend where the control group had a higher daily consumption (Fig. 3A). Then, both groups stabilized their consumption at about 31 g/day (Fig. 3A). Statistical analysis revealed a significant effect of time (F (3.440, 75.69) = 38.73, P<0.0001; Greenhouse–Geisser ε = 0.49), no effect of diet (F (1, 22) = 0.3539, P=0.5580) and a significant time X diet interaction (F (7, 154) = 15.95, P<0.0001). In adult female rats, the HFSD group had a higher consumption of food than the control group for the first 5 weekends (20.0 ± 0.2 g/day for the control group and 21.8 ± 0.3 g/day for the HFSD group) (Fig. 3B). Then, the consumption stabilized and became similar in two groups at about 19 g/day (Fig. 3B). Statistical analysis on daily consumption revealed a significant effect of time (F (3.194, 70.26) = 20.53, P<0.0001; Greenhouse–Geisser ε = 0.46), a significant effect of diet (F (1, 22) = 11.21, P=0.0029) and a significant time X diet interaction (F (7, 154) = 4.108, P=0.0004). In adolescent male rats, during the first 2 weekends, the HFSD group had a lower daily consumption than the control group (∼25 g/day for the control group ∼23 g/day for the HFSD group) then, on the last weekend, the consumption of HFSD group was higher than control group (∼31g/day for the control group and ∼34g/day for the HFSD group) (Fig. 3C). Statistical analysis revealed a significant effect of time (F (3.762, 82.77) = 235.1, P<0.0001; Greenhouse–Geisser ε = 0.54), no significant effect of diet (F (1, 22) = 0.8806, P=0.3582) and a significant time X diet interaction (F (7, 154) = 11.88, P<0.0001). In adolescent female rats, the HFSD and control group initially had a similar daily consumption of food but after 3-4 weeks HFSD rats started consuming more food than controls (19.5 ± 0.4 g/day for the control group and 21.2 ± 0.5 g/day for the HFSD group) (Fig. 3D).

**Fig 3:**
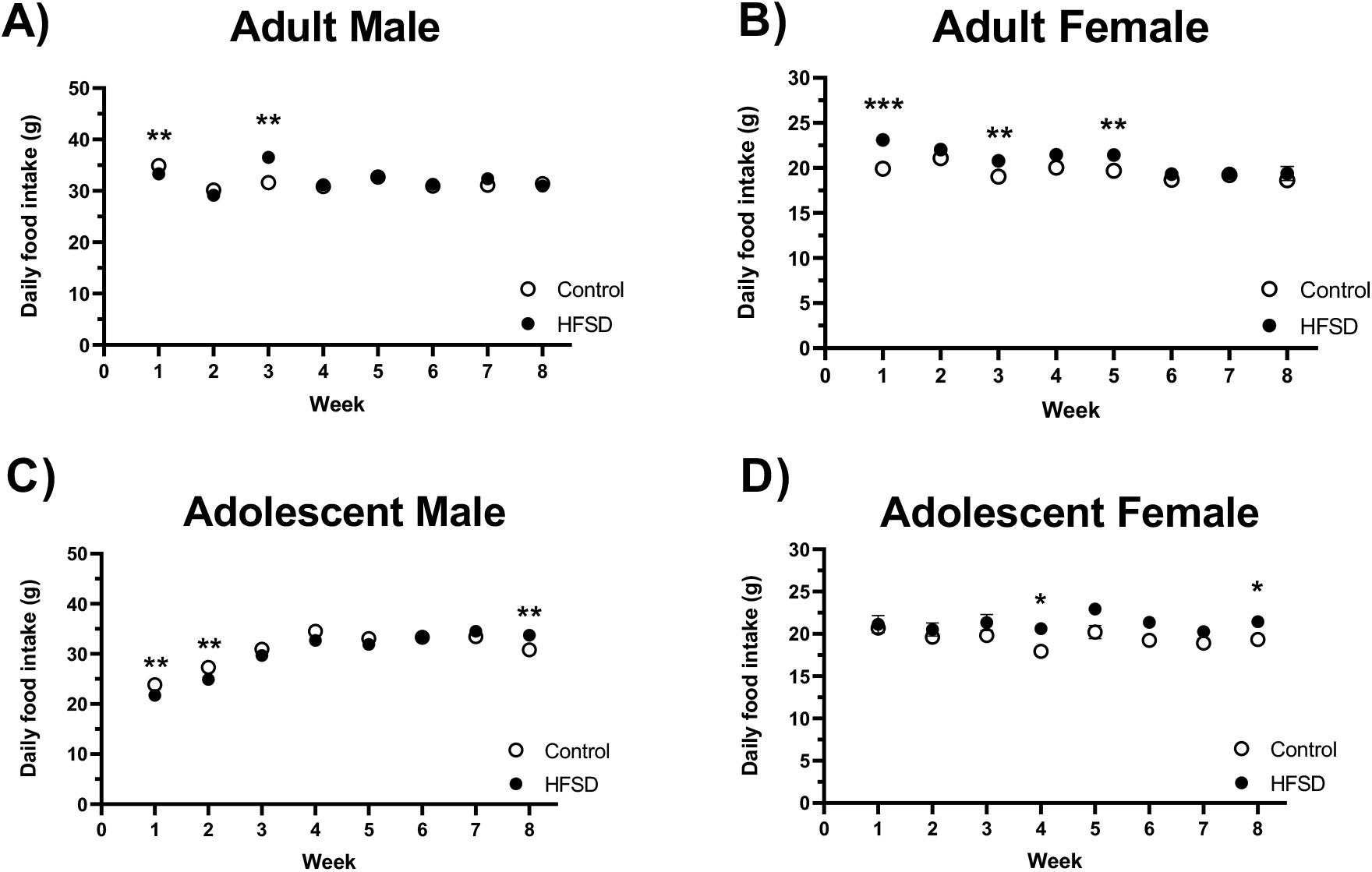
Effects of HFSD on daily food consumption during weekends. Time course of daily food consumption in the weekends during the 8 weeks of exposure to HFSD in rats exposed to standard food (Control) and rats exposed to HFSD (HFSD). A) Adult males (HFSD n=12; Control n=12); B) Adult females (HFSD n=12; Control n=12); C) Adolescent males (HFSD n=12; Control n=12) and D) Adolescent females (HFSD n=12; Control n=11). Data are expressed as Mean ± SEM. * P<0.05; ** P<0.01; *** P<0.001

Statistical analysis revealed a significant effect of time (F (3.322, 69.77) = 5,515, P=0.0013; Greenhouse–Geisser ε = 0.47), a significant effect of diet (F (1, 21) = 8.067, P=0.0098) and no significant time X diet interaction (F (7, 147) = 1.772, P=0.0971).

Concerning daily food consumption during the rest of the week, when all groups had access only to standard food, in adult male rats, the control group had a higher consumption than the HFSD group during weekdays (∼31g/day for the control group and ∼27 g/day for the HFSD group) (Fig. 4A). Statistical analysis revealed a significant effect of time (F (2.987, 65.72) = 24.18, P<0.0001; Greenhouse–Geisser ε = 0.43), a significant effect of diet (F (1, 22) = 15.38, P=0.0007) and a significant time X diet interaction (F (7, 154) = 7.371, P<0.0001). In adult female rats, the HFSD group had a lower daily consumption than the control group (18.7 ± 0.3 g/day for the control group and 16.3 ± 0.2g/day for the HFSD group) (Fig. 4B). Statistical analysis revealed a significant effect of time (F (1.768, 38.90) = 8.768, P=0.0011; Greenhouse–Geisser ε = 0.25), a significant effect of diet (F (1, 22) = 39,70, P<0.0001) and a significant time X diet interaction (F (7, 154) = 2.133, P=0.0433). In adolescent male rats, HFSD and control group initially had similar consumption but in the last 4 weeks the consumption of HFSD rats was lower than the control (32.2 ± 0.2 g/day for the control group and 29.6 ± 0.2 g/day for the HFSD group) (Fig. 4C). Statistical analysis revealed a significant effect of time (F (1.545, 34.00) = 77.16, P<0.0001; Greenhouse–Geisser ε = 0.22) a significant effect of diet (F (1, 22) = 32.98, P<0.0001) and a significant time X diet interaction (F (7, 154) = 10.42, P<0.0001). In adolescent female rats, the HFSD group had a lower consumption of food the control (21.5 ± 0.3 g/day for the control group and 20.0 ± 0.4 g/day for the HFSD group about the first weekend and 19.7 ± 0.4 g/day for the control group and 18.6 ± 0.3 g/day for the HFSD group on the last weekend) (Fig. 4D). Statistical analysis revealed a significant effect of time (F (3.951, 82.98) = 16.38, P<0.0001; Greenhouse–Geisser ε = 0.56), a significant effect of diet F (1, 21) = 5.383, P=0.0305) and a significant time X diet interaction (F (7, 147) = 3.055, P=0.0050).

**Fig 4:**
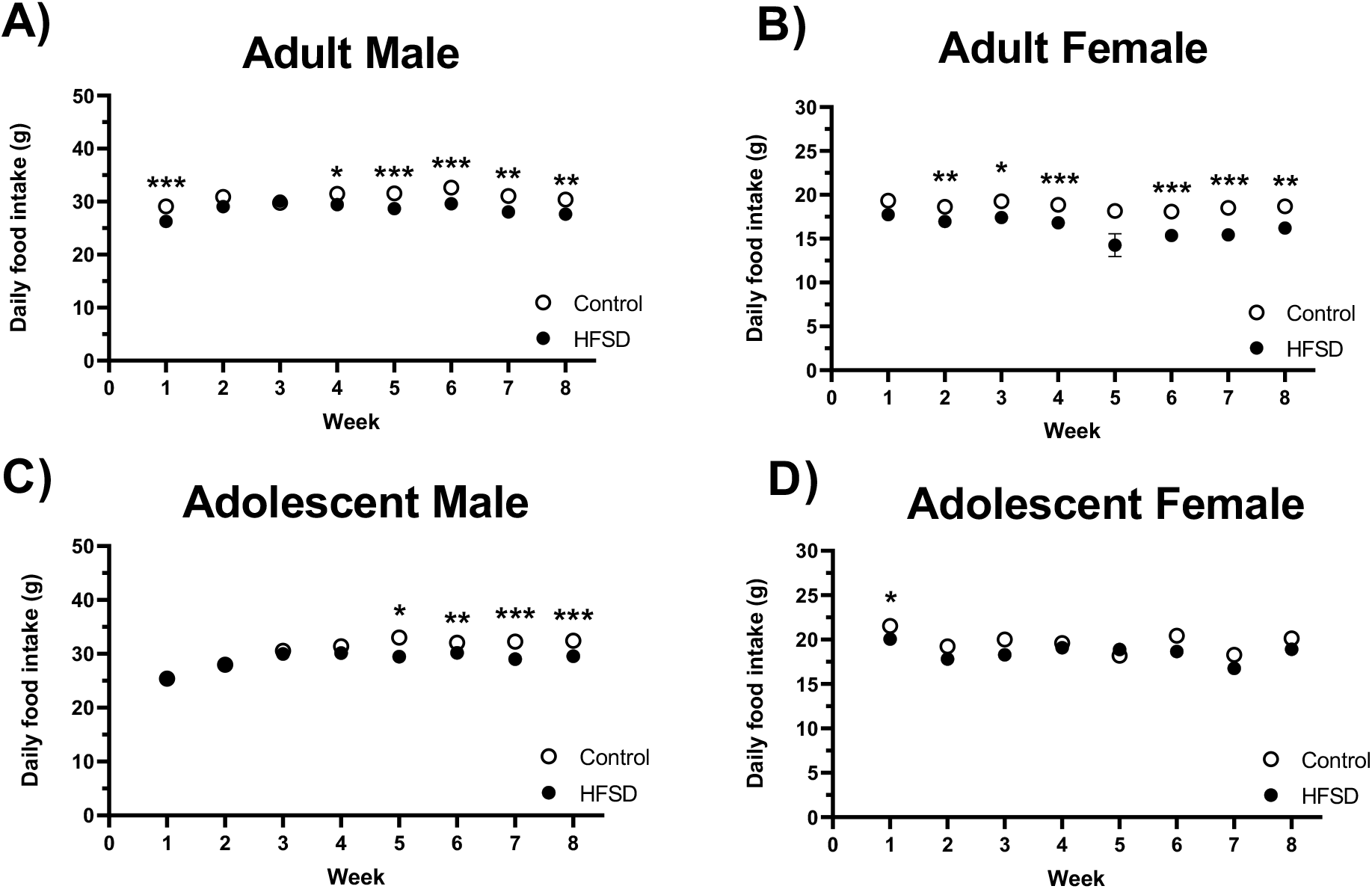
Effects of HFSD on daily food consumption during weeks. Time course of daily food consumption in the week during the 8 weeks of exposure to HFSD in rats exposed to standard food (Control) and rats exposed to HFSD (HFSD). It should be noted that only standard food was available during the week for both HFSD and controls. A) Adult males (HFSD n=12; Control n=12); B) Adult females (HFSD n=12; Control n=12); C) Adolescent males (HFSD n=12; Control n=12) and D) Adolescent females (HFSD n=12; Control n=11). Data are expressed as Mean ± SEM. * P<0.05; ** P<0.01; *** P<0.001

In summary, exposure to intermittent HFSD did not result in increases in body weight in any group. When rats had access to HFSD, they tended to consume less food compared to animals that had access to standard food. More importantly, exposure to HFSD reduced consumption of standard food during the weekdays suggesting that exposure to HFSD made standard food less rewarding.

#### Effects of exposure HFSD on food taking in the FR1

During the first 5 training sessions in the FR1, we found no difference between HFSD and control rats in any condition (fig. S1) suggesting that operant learning was not affected by HFSD.

In contrast, in adult male rats, baseline responding in the FR1 schedule (the average responding in the 3 sessions before PSS tests) was lower in HFSD rats compared to controls in the first weeks after discontinuation of HFSD and then reached control levels in the last 2 weeks (Fig.5A). Statistical analysis revealed a significant effect of time (F (2.270, 49.93) = 40.39; P<0.0001; Greenhouse–Geisser ε = 0.57), a significant effect of diet (F (1, 22) = 14.90; P = 0.0008) and a significant Time*Diet interaction (F (4, 88) = 6.181 P = 0.0002). Similarly, in adult female rats, FR1 baseline responding was lower in the HFSD early after discontinuation of HFSD and then returned to control levels (Fig.5B). Statistical analysis revealed a significant effect of time (F (1.950, 42.90) = 55.40, P < 0.0001; Greenhouse–Geisser ε = 0.49), significant effect of diet (F (1, 22) = 5,263, P = 0.037) but not significant Time*Diet interaction. In adolescent male rats, initial baseline responding was slightly lower after discontinuation and rapidly returned to control levels (Fig.5C). Statistical analysis revealed a significant effect of time (F (1.659, 36.51) = 50.58, P < 0.0001; Greenhouse–Geisser ε = 0.42), no significant effect of diet but a significant Time*Diet interaction (F (4, 88) = 3.581; P=0.0094). No difference in baseline responding in the FR1 schedule was found in female adolescent rats (Fig.5C) and statistical analysis revealed only a significant time effect (F (3.345, 70.25) = 5.906; P=0.0008; Greenhouse–Geisser ε = 0.83)

**Fig 5:**
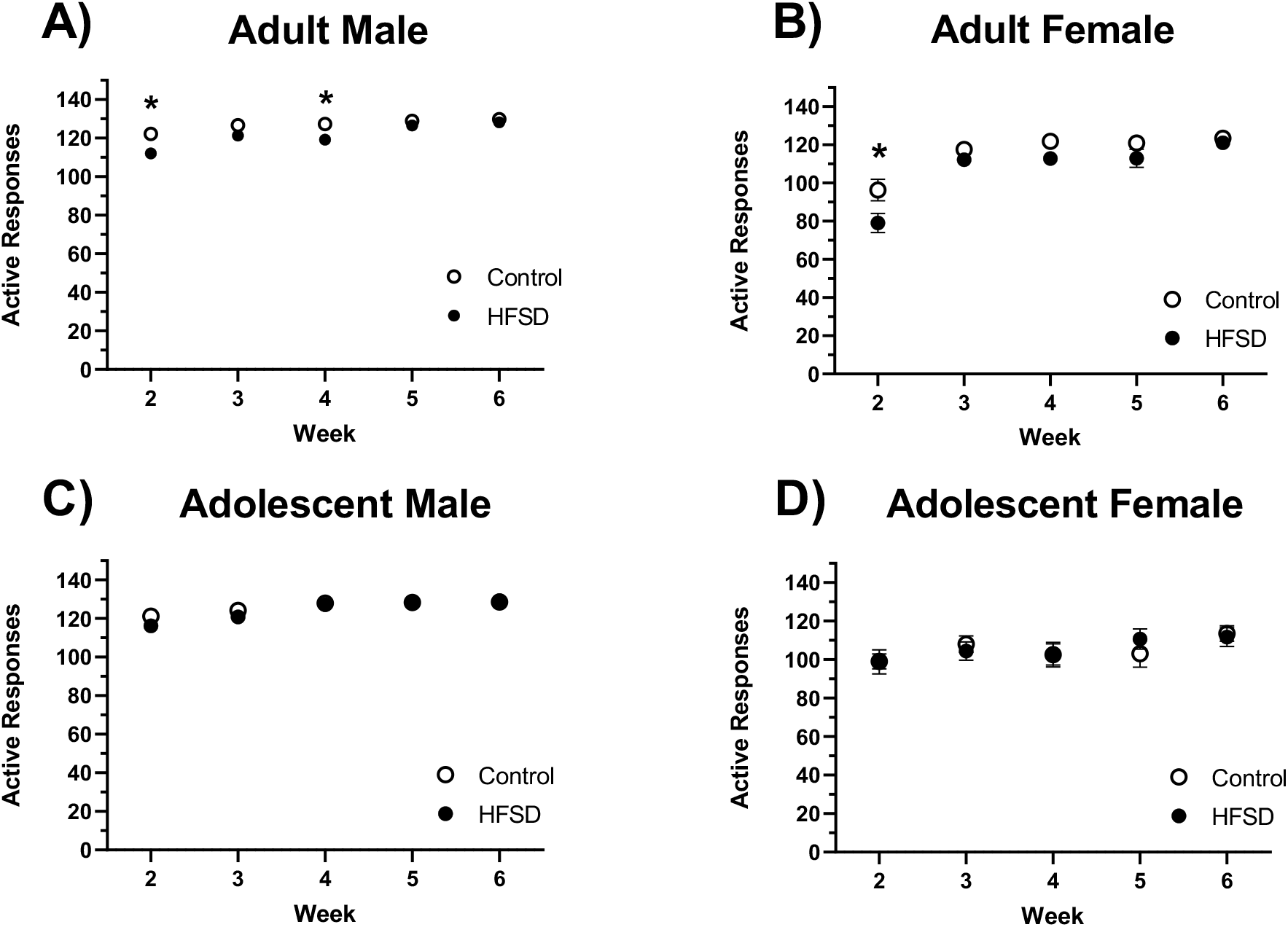
Effects of HFSD on baselines in the training FR1 sessions during the experiment. Time course the baseline during the experiments in rats exposed to standard food (Control) and rats exposed to HFSD (HFSD). It should be noticed that the first week of training in which learning occurred is not represented here. It should also be noticed that food was discontinued about 7 days before the start of FR1 training. A) Adult males (HFSD n=12; Control n=12); B) Adult females (HFSD n=12; Control n=12); C) Adolescent males (HFSD n=12; Control n=12) and D) Adolescent females (HFSD n=12; Control n=11). Data are expressed as Mean ± SEM. * P<0.05.

#### Exposure to HFSD does not produce a persistent alteration in compulsive food taking

In adult males (Fig. 6A) and female rats (Fig 6B) and in adolescent males (Fig 6C) and female rats (Fig 6D), HFSD and control groups had a similar PSS break point (adult males: 1,35 ± 0,5 mC for the control group and 0,80 ± 0,3 for the HFSD group (P = 0.09; t = 1.77; DF = 22); adult females: 1.6 ± 0.4 mC for the control group and 1.4 ± 0,5 (P = 0.40; t = 0.85; DF = 22); adolescent males: 1.1 ± 0.2 mC for the control group and 0.8 ± 0,1 (P = 0.20; t = 1.331; DF = 22); adolescent females: 0.5 ± 0.1 mC for the control group and 0.8 ± 0,1 (P = 0.87; t = 0.16; DF = 21).

**Fig 6:**
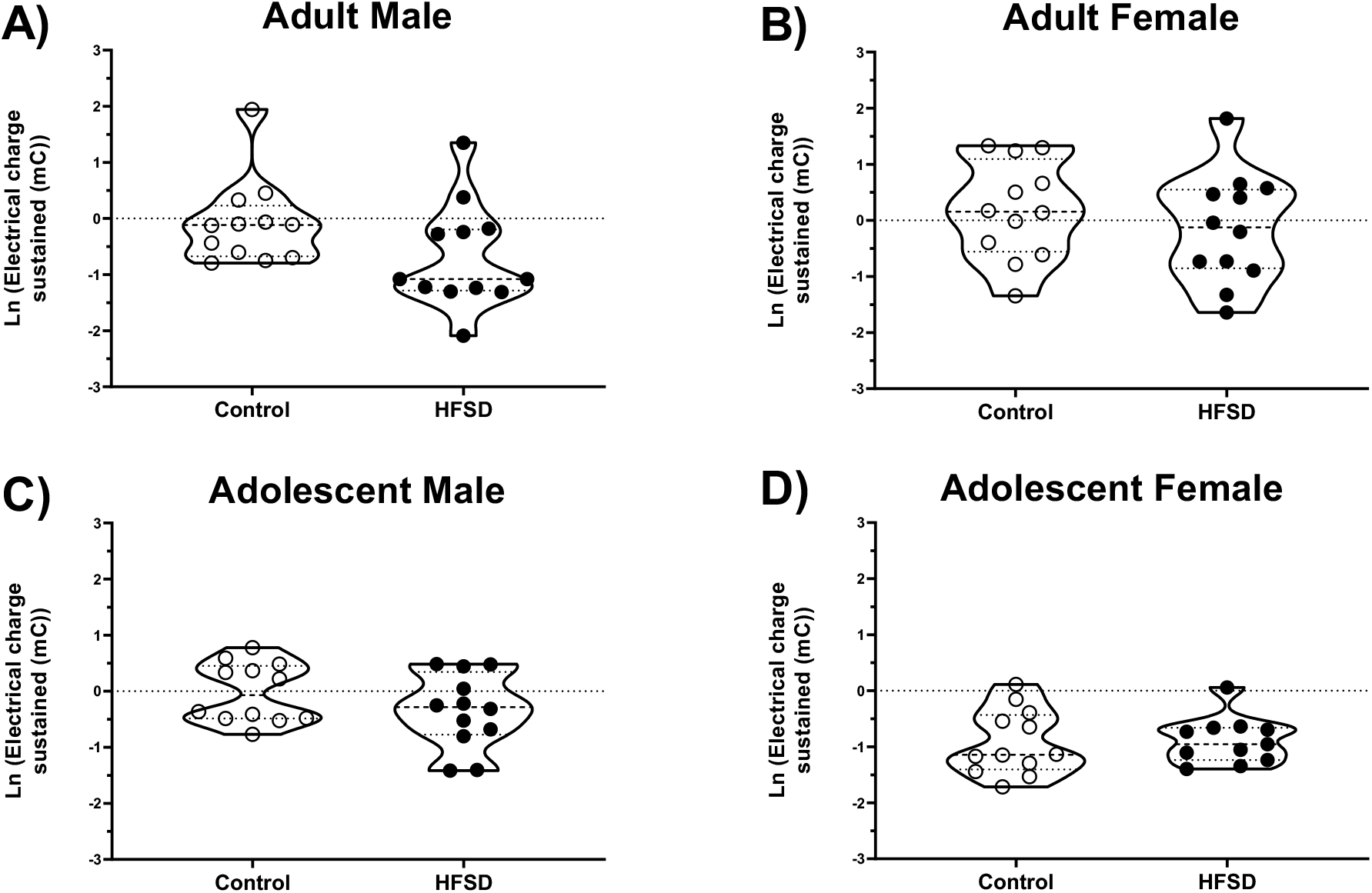
Effect of HFSD on resistance to punishment in the PSS procedures. Natural logarithm of electric charge sustained in the PSS tests in rats exposed to standard food (Control) and rats exposed to HFSD (HFSD). A) Adult males (HFSD n=12; Control n=12); B) Adult females (HFSD n=12; Control n=12); C) Adolescent males (HFSD n=12; Control n=12) and D) Adolescent females (HFSD n=12; Control n=11). Data represent the average of two sessions and are expressed as Mean ± SEM.

#### Exposure to HFSD does not produce persistent alterations in the motivation for food

In adult males (Fig. 7A) and female rats (Fig 7B) and in adolescent males (Fig 7C) and female rats (Fig 7D), HFSD and control groups had a similar PR break point (adult males: 581 ± 45 for the control group and 480 ± 42 active responses for the HFSD group (P = 0.12; t = 1.64; DF = 22) adult females: 413 ± 64 for the control group and 407 ± 45 active responses for the HFSD group (P = 0.93; t = 0.08; DF = 22); adolescent males: 397 ± 38 for the control group and 380 ± 45 active responses for the HFSD group (P = 0.77; t = 0.30; DF = 22); adolescent females: 236 ± 32 for the control group and 188 ± 35 active responses for the HFSD group (P = 0.34; t = 0.99; DF = 21).

**Fig 7:**
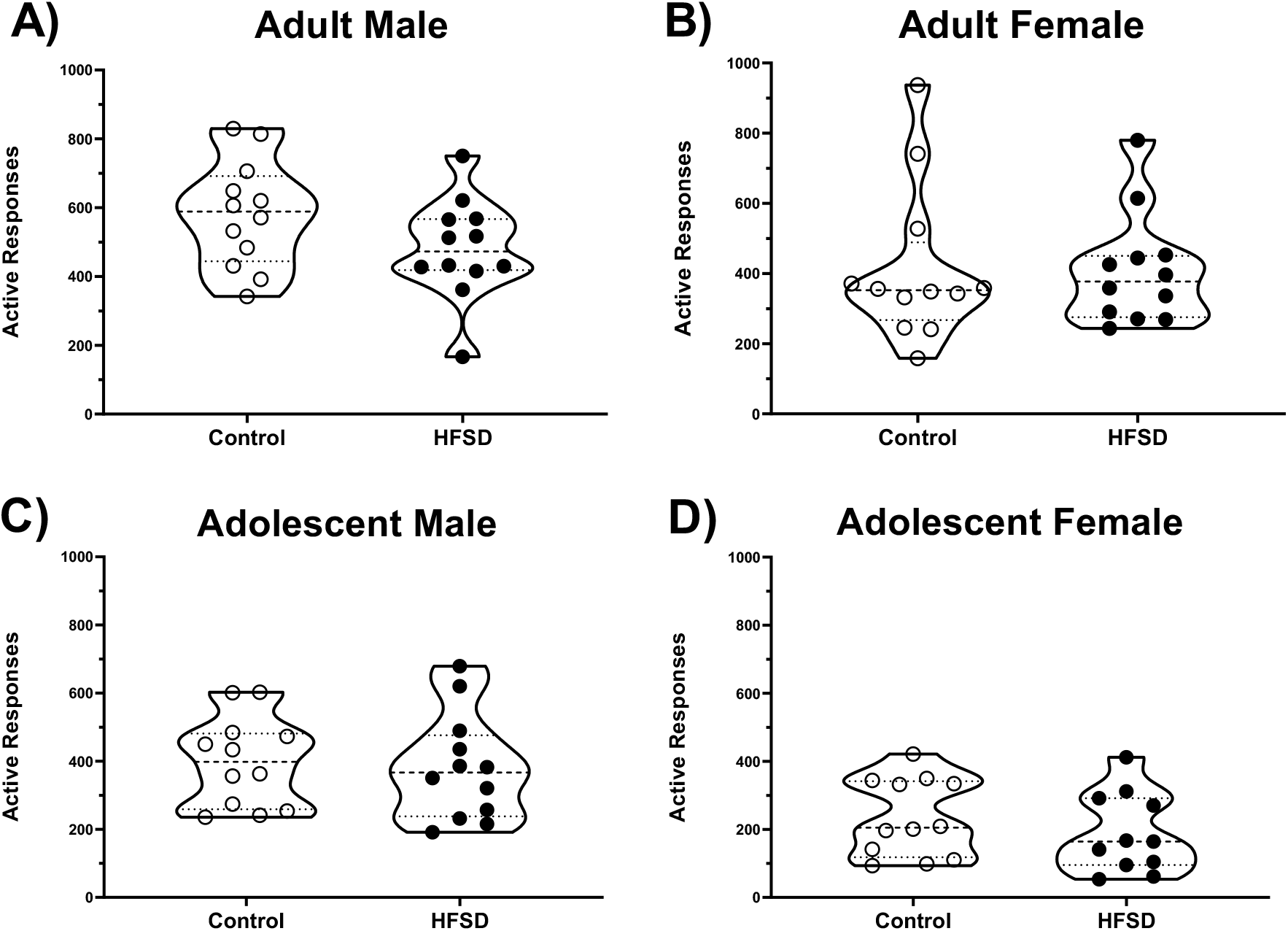
Effect of HFSD on food motivation in the PR procedure. Total active responses in the PR tests in rats exposed to standard food (Control) and rats exposed to HFSD (HFSD). A) Adult males (HFSD n=12; Control n=12); B) Adult females (HFSD n=12; Control n=12); C) Adolescent males (HFSD n=12; Control n=12) and D) Adolescent females (HFSD n=12; Control n=11). Data represent the average of two sessions and are expressed as Mean ± SEM.

#### Effects of exposure to HFSD on anxiety-like behaviors and pain sensitivity

For anxiety-like behavior, HFSD and control groups spent similar amount of in the open arms regardless of sex or age. In adult male rats, HFSD spent 88 ± 54 s in open arms compared to the control group with 94 ± 31 s (Fig. 8A). Statistical analysis on time spent in open arms revealed no significant effect of diet (Mann-Whitney U = 38.50; P = 0.053). In adult female rats, HFSD spent 77 ± 20 s in open arms compared to the control group with 115 ± 33 s (Fig. 8B) Statistical analysis on time spent in open arms revealed no significant effect of diet (Mann-Whitney U = 59; P = 0.48). In adolescent male rats, HFSD spent 54 ± 12 s in open arms compared to the control group with 58 ± 12 s (Fig. 8C). Statistical analysis on time spent in open arms revealed no significant effect of diet (Mann-Whitney U = 69.50; P = 0.90). In adolescent female rats, HFSD spent 107 ± 14 s compared to the control group with 142 ± 25 s (Fig. 8D). Statistical analysis on time spent in open arms revealed no significant effect of diet (Mann-Whitney U = 48, P = 0.29).

**Fig 8:**
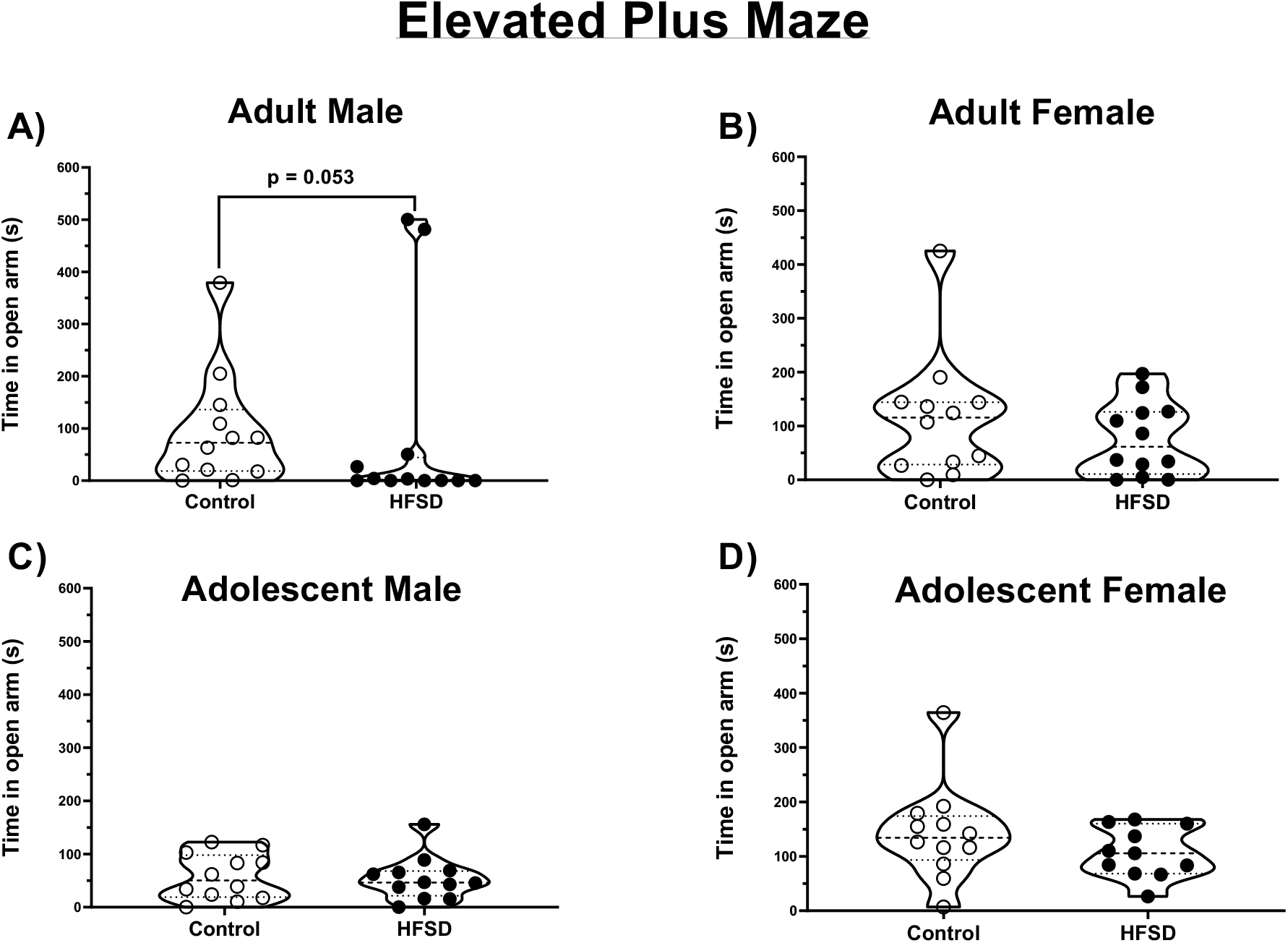
Effects of HFSD on anxiety-like behavior. Time spent in second in the open arms of an elevated plus maze in rats exposed to standard food (Control) and rats exposed to HFSD (HFSD). A) Adult males (HFSD n=12 ; Control n=12); B) Adult females (HFSD n=12 ; Control n=12); C) Adolescent males (HFSD n=12 ; Control n=12) and D) Adolescent females (HFSD n=12 ; Control n=11). Data are expressed as Mean ± SEM.

In the hot plate test, in adult males, HFSD and control groups had similar latency before to trying to escape (93 ± 6s for the control group and 91 ± 9 s for the HFSD group) (Fig. 9A). Statistical analysis on the latency before escape revealed no significant effect of diet (Mann-Whitney U = 67; P = 0.78). In adult females, HFSD and control groups had similar latency before to trying to escape (100 ± 9 s for the control group and 92 ± 7 s for the HFSD group) (Fig. 9B). Statistical analysis on the latency before escape revealed no significant effect of diet (Mann-Whitney U = 56; P = 0.35). In adolescent males, the latency was lower in HFSD compared to the control (109 ± 9 s for the control group and 84 ± 9 s for the HFSD group) suggesting that in male exposure to HFSD during adolescence produces long-lasting increases in pain sensitivity (Fig. 9C). Statistical analysis on the latency before escape revealed a significant effect of diet (Mann-Whitney U = 36; P=0.03). In adolescent females, HFSD and control groups had similar latency before to trying to escape (79 ± 13 s for the control group and 67 ± 14 s for the HFSD group) (Fig. 9D). Statistical analysis on the latency before escape revealed a significant effect of diet (Mann-Whitney U = 49; P=0.30).

**Fig 9:**
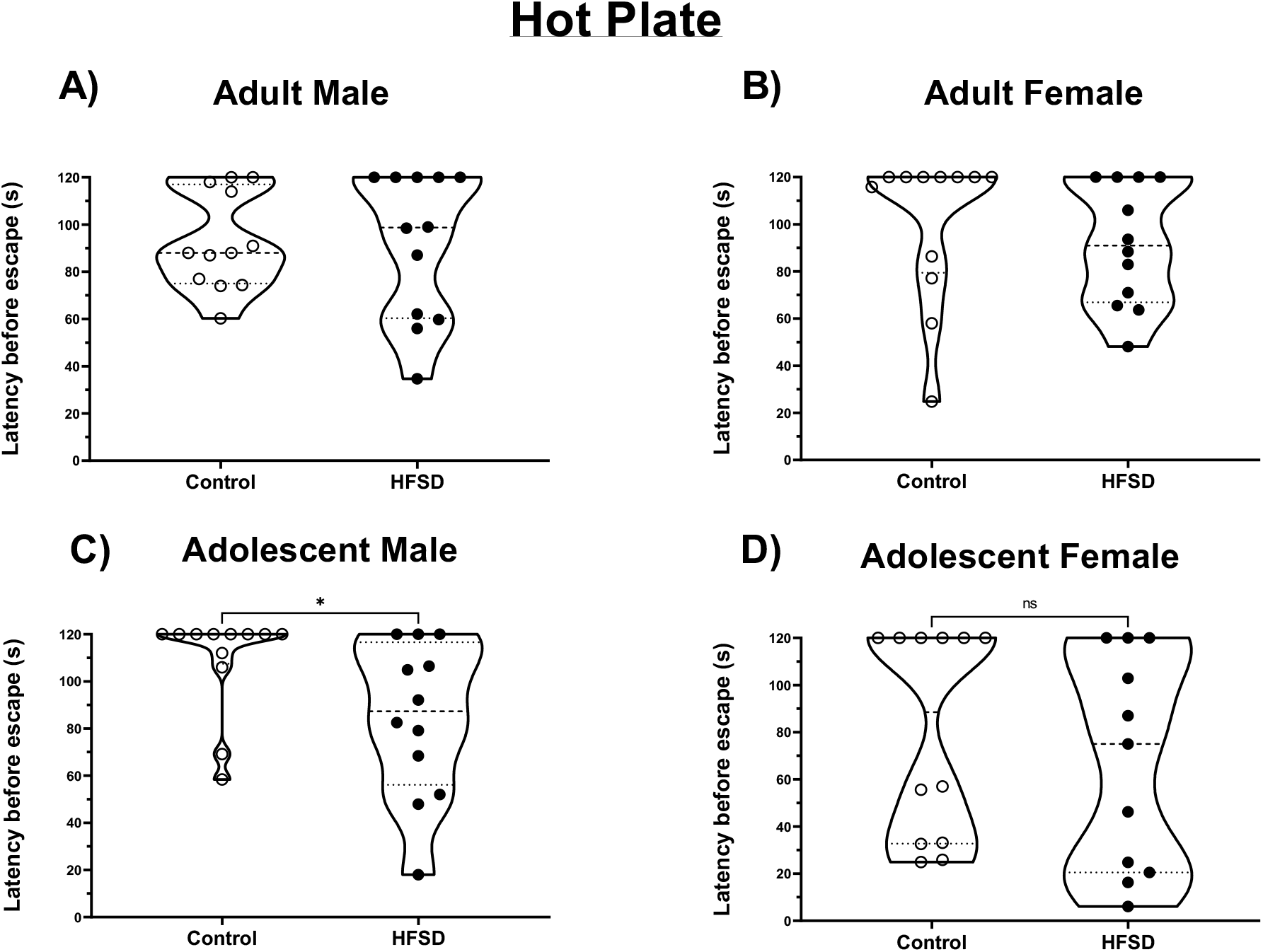
Effects of HFSD on pain reactivity by hot plate test. Latency in second before escape in the hot plate test in rats exposed to standard food (Control) and rats exposed to HFSD (HFSD). A) Adult males (HFSD n=12 ; Control n=12); B) Adult females (HFSD n=12; Control n=12); C) Adolescent males (HFSD n=12 ; Control n=12) and D) Adolescent females (HFSD n=12; Control n=11). Data are expressed as Mean ± SEM. * P<0.05.

## DISCUSSION

In this study, we investigated the effects of intermittent exposure to HFSD on subsequent food self-administration and notably, on indexes of compulsivity and excessive motivation. We found that, irrespectively of sex or age, when rats had access to HFSD, their pattern of food intake was altered with decrease of consumption of standard chow in between periods of HFSD exposure. In addition, in adult but not adolescent rats, exposure to HFSD produced a transient decrease in self-administration of sucrose under FR1 schedule. However, HFSD did not produce persistent alterations in compulsivity or motivation for food which would suggest an increase in the risk to develop addiction-like behavior towards food.

Exposure to highly processed food rich in fat and sugar is believed to be a major risk factor to develop excessive eating and overweight and obesity (O’Connor and Kenny, 2022). In our study, intermittent exposure to HFSD diet did not induce increase in body weight suggesting that this type of exposure is not necessarily obesogenic. This is consistent with previous reports showing no increase in body weight in animals exposed intermittently to HFSD (Cottone et al., 2009; Ferragud et al., 2020; Rossetti et al., 2014). On the other hand, exposure to HFSD altered the pattern of food intake during the period of exposure in all groups. In particular, consistent with previous studies (Cottone et al., 2009; Ferragud et al., 2020; Rossetti et al., 2014), HFSD rats showed a decrease in consumption of standard food during the week suggesting a negative contrast effect with exposure to high palatable food decreases the rewarding of effect of normal food. Female rats (adult and adolescent), consumed more HFSD food compared to control groups confirming that, in females, HFSD is more rewarding than standard food. In contrast, adult and adolescent male rats showed mixed results in HFSD consumption with low initial consumption, probably due to novelty avoidance, followed by a slight increase in consumption in the subsequent weeks. Thus, male rats appear less sensitive to the rewarding effects of HFSD than female rats.

Resistance to punishment is often used as index of compulsivity for food or drugs. In many procedures in rodents, punishment is provided as a foot shock of a fixed intensity that allows characterizing animals in resistant (i.e. compulsive) and sensitive (i.e. non-compulsive) (George et al., 2022; Rossetti et al., 2014; Vanderschuren and Ahmed, 2013). However, using an arbitrary threshold forces a dichotomic classification ignoring the fact that compulsivity may be a more continuous phenomenon. To avoid this problem and to better mimic tests of compulsivity used in humans (Apergis-Schoute et al., 2017; Kanen et al., 2021; Kim and Anderson, 2020), we recently developed a self-adjusting procedure that allows individual animals to titrate the level of shock that they are willing to tolerate to obtain rewards (Desmercieres et al., 2022). Consistent with our previous study, we found that compulsivity measured by the PSS break point is log-normally distributed suggesting that resistance to punishment is a continuous rather than dichotomic phenomenon regardless of sex or exposure to HFSD. In addition, we found that regardless of sex or age of HFSD exposure PSS and PR do not correlate (Supplementary Fig. 3) confirming that they measure independent biobehavioral processes.

Previous studies have found that exposure to high fat and sugar diet increases (Garman et al., 2021; la Fleur et al., 2007; Reichelt et al., 2016; Wojnicki et al., 2006), decreases (Blaisdell et al., 2014; Davis et al., 2008; Tracy et al., 2015; Vendruscolo et al., 2010) or has no effect (Jong et al., 2013; Mitra et al., 2009; Rossetti et al., 2014) on motivation for food measured by progressive ratio schedules. Some of these discrepancies could be due to the sex of animals since most studies investigated males and females separately. One study that directly investigated male and female rats exposed to sucrose found that increases motivation in female but it decreases in male rats (Reichelt et al., 2016). Differently from that study, we found that HFSD did not change motivation for sweet sucrose rewards in males and females exposed to HFSD as adults and adolescents. Another possible reason for discrepancies found in previous studies is the age at which animals were exposure to obesogenic diets because most studies investigated the effects of exposure at adolescence or adulthood separately. Indeed, adolescence is a window of vulnerability for the development of maladaptive behaviors including addiction food-related disorders (Larsen and Luna, 2018; Murray and Chen, 2019) and it has been shown that exposure to obesogenic diets at adolescence leads to increased (Figlewicz et al., 2013) or decreased (Vendrusdcolo et al., 2010) motivation compared to exposure at adulthood. However, in our study, motivation for sweet food was equally unaffected by exposure during adolescence or adulthood.

Concerning compulsivity measured as resistance to punishment, the results in the literature also show contrasting results with some studies reporting increases (Oswald et al., 2011; Rossetti et al., 2014) and another no change (Jong et al., 2013) after exposure to obesogenic diets. In addition, Johnson and Kenny found that prolonged access to an obesogenic diet blunted the ability of an aversive conditioned stimulus to disrupt food seeking and taking (Johnson and Kenny, 2010). In the present study, using the PSS procedure we found no effect of exposure to HFSD on compulsivity regardless of sex and age of exposure to HFSD suggesting that these factors may not explain by themselves the differences in the literature.

In this study, we wanted to investigate the long-lasting rather than the immediate consequences of exposure to obesogenic diets because if diet-induced dysfunctions persist even when bad eating habits are changed for healthier ones, this would have more serious consequence on health. For this reason, after 8 weeks of exposure to HFSD, rats were switched to normal chow for several days before being tested for excessive motivation or resistance to punishment. This may have contributed to the lack of effects of HFSD since other studies have found that some of the deleterious effects of obesogenic diets tend to disappear upon discontinuation (Boitard et al., 2016; Carlin et al., 2016; Rabasa et al., 2016). However, other studies have found long-lasting effects of obesogenic diets on motivation (Garman et al., 2021; Reichelt et al., 2016; Tracy et al., 2015; Vendruscolo et al., 2010). Our protocol of exposure was based on the one used by Rossetti et al. (2014) except that we used Sprague-Dawley instead of Wistar rats and we discontinued HFSD diet several days before the beginning of operant training. Whereas Rossetti et al. found increases in resistance to punishment in females exposed to HFSD at adolescence (Rossetti et al., 2014), we did not find significant difference in any of our comparisons. Future studies are needed to determine whether these discrepancies are due to rats’ strain or discontinuation of HFSD exposure.

In humans, obesity is often associated with anxiety (Fulton et al., 2022; Gariepy et al., 2010) but the causal relationship between the two conditions remains unclear. Some studies in rats have shown that obesogenic diets induce increases in some tests of anxiety (Cottone 2009 ; Sivanathan et al. 2015; Dutheil et al. 2016) but little or no effect has been described in the elevated plus maze (Dutheil et al., 2016; Rossetti et al., 2014; Sivanathan et al., 2015). Consistent with these latter findings, we found no significant effect of HFSD on anxiety in the elevated plus maze in any group although there was a general trend for increased anxiety in all groups and the trend almost reached significance in the adult male groups. It should be considered that anxiety was only evaluated at the end of our food self-administration, several weeks after exposure to HFSD. Therefore, we cannot exclude exposure to HFSD induced anxiety at earlier time point, which disappeared over time.

In humans, obesity is also associated with pain and the relationship appears bidirectional (McVinnie, 2013). According to a metanalysis of the field, experiments in rodents mostly found a decrease in pain sensitivity in a variety of conditions and tests (Marques Miranda et al., 2021). However, some studies have also found increased pain sensitivity (Roane and Porter, 1986; Song et al., 2017; Tramullas et al., 2016) and this may be due to inflammatory processes (Song et al., 2017; Tramullas et al., 2016). In our study, consistent with these latter studies, we found a general trend for increased sensitivity in all groups that reached statistical significance in adolescent male rats. Again, pain sensitivity was measured at the end of the experiments and therefore, it is possible that HFSD produce a temporary increase in pain sensitivity which tends to recover upon discontinuation of HFSD. Interestingly, if pain sensitivity was increased by exposure to HFSD during the first days/weeks of withdrawal, this did not affect PSS breakpoints. This confirm previous studies (Degoulet et al., 2021; Desmercieres et al., 2022; Li et al., 2021) showing that pain sensitivity does not play a major role in resistance to punishment.

Important issues in the investigation of the effects of HFSD on brain and behavior are both the actual composition of diets and the protocol of exposure. Indeed, some studies, similar to ours, used enriched tablets (Cottone et al., 2009; Iemolo et al., 2012; Rossetti et al., 2014), others used sucrose solutions (Reichelt et al., 2016; Vendruscolo et al., 2010; Wong et al., 2017) and others different kinds of commercial or home-made sugary and fat diets (Johnson and Kenny, 2010; Jong et al., 2013; la Fleur et al., 2007; Oswald et al., 2011; Tracy et al., 2015). In addition, the duration of the exposure varies from a few days (Oswald et al., 2011), to several weeks and up to six months (Blaisdell et al., 2014). Finally, the timing of the exposure could be continuous (Blaisdell et al., 2014; Davis et al., 2008; Figlewicz et al., 2013; Garman et al., 2021; la Fleur et al., 2007; Tracy et al., 2015; Vendruscolo et al., 2010), intermittent with periods of several days (Cottone et al., 2009; Iemolo et al., 2012; Johnson and Kenny, 2010; Rossetti et al., 2014) or a few hours per day (Reichelt et al., 2016; Wojnicki et al., 2006; Wong et al., 2017). It is difficult to determine which procedure better mimics human exposure to HFSD because human food selection and intake varies enormously not only in different cultures and in different socio-economical classes, but also within them and in many instances even within the same person at different times. Whereas all these differences in experimental protocols are likely to account for the discrepancies in the literature, we should probably embrace the richness provided by this variety because findings that are consistent among different procedures are likely to have the highest translational value for human condition.

In conclusion, this study found that intermittent exposure to HFSD does not produce persistent effects on motivation and compulsivity to consume sweet food regardless of sex or age of exposure. This lack of effects, together with the great number of contradicting results found in the literature suggest that the persistent consequences of exposure to HFSD on the development of food addiction are at best limited and subtle.

