## Supplementary figures and images for "Effects of high fat and sugar diet on motivation for food and resistance to punishment in rats: role of sex and age of exposure"

### Fig. S1

#### Slide 1
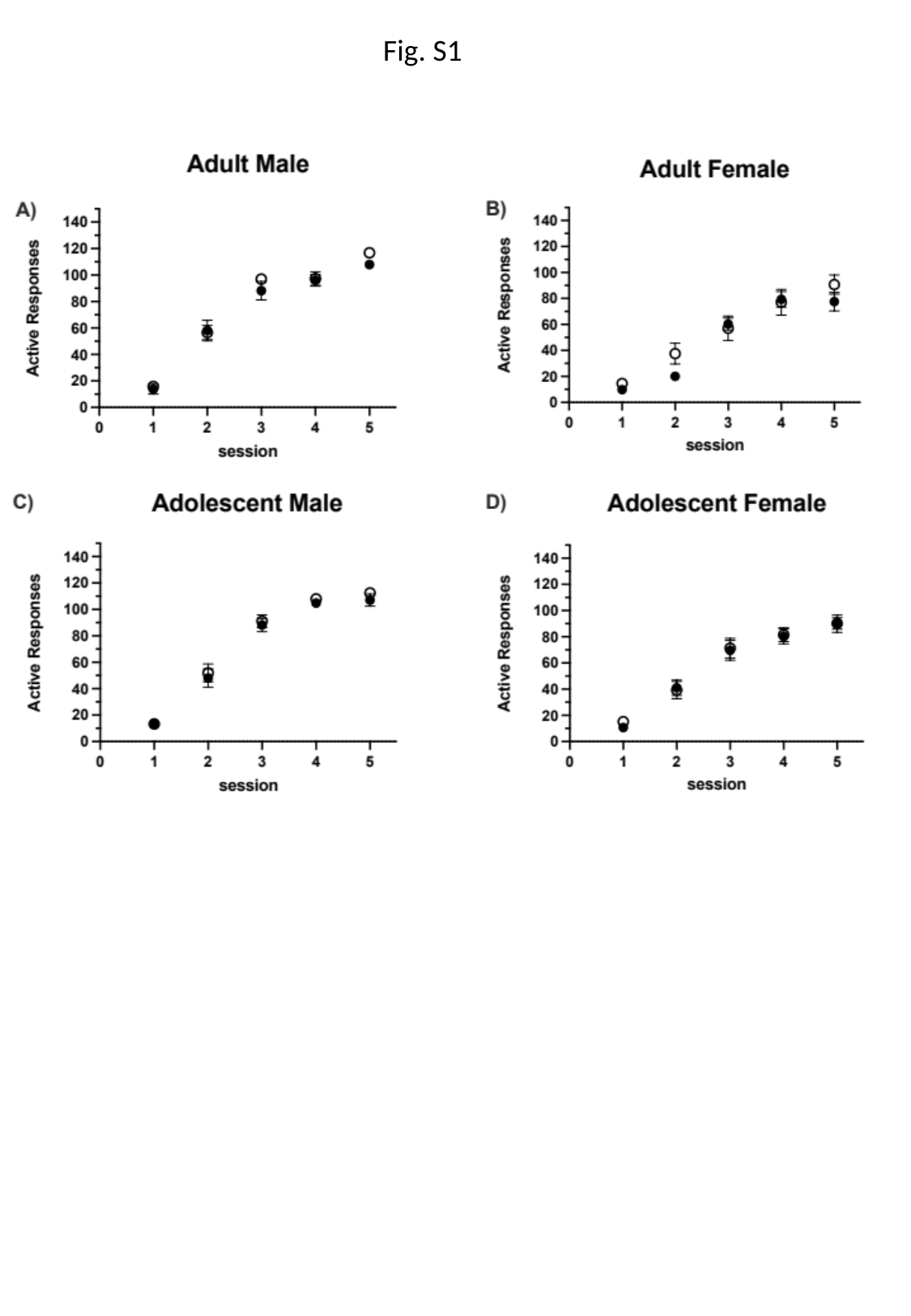

Fig. S1
